# Oscillatory Multi-Timescale Mechanisms Underlying Audiovisual Sequence Prediction

**DOI:** 10.1101/778969

**Authors:** Peng Wang, Alexander Maÿe, Jonathan Daume, Gui Xue, Andreas K. Engel

## Abstract

Neural oscillations have been proposed to be involved in predictive processing, however less addressed in the multisensory context. In the present study, we recorded cortical activity using magnetoencephalography during an audiovisual serial prediction task in which participants had to acquire stimulus sequences and to monitor whether subsequent probe items complied with the sequence. In a given trial, either visual or auditory features of the bimodal stimuli were task-relevant. Task-related changes of power and coupling of neural oscillations were analyzed using a data-driven clustering approach. We observed prediction-related theta-band (5-7 Hz) coupling in a network involving the cingulate, premotor, prefrontal, and superior temporal regions. The behavioral performance of participants varied as a function of the phase delay of the coupling between different regions in this network. . Two additional spectro-temporal clusters were detected in which beta power (11-16 Hz and 16-22 Hz, respectively) was stronger if the previous or the following stimuli were the same as the current one in the sequence. These clusters spatially overlapped with the theta-band prediction cluster, and interacted with it through cross-frequency coupling. Our results offer novel insights into the multi-timescale neural dynamics underlying multisensory predictions, suggesting that oscillations in multiple frequency ranges, as well as coupling within and across frequencies, may be critical for multisensory sequence processing.

## 1 Introduction

Natural events and objects are often jointly detected by more than one sensory system. Their perceptual and behavioral consequences are often not just the sum of those evoked by each sensory component alone. However, our knowledge is still limited regarding the mechanisms how multisensory interactions in the human brain contribute to unified perceptual experiences and decisions, reviewed in (Bauer et al., 2020; Senkowski & Engel, 2024; Senkowski et al., 2008; Talsma et al., 2010). Fewer studies have so far addressed the neural substrates of multisensory processing with data-driven measures of large-scale neural dynamics (Cooke et al., 2019; Galindo-Leon et al., 2019; Hipp et al., 2011; Keil et al., 2012; Wang et al., 2019). Recent electroencephalography (EEG) and magnetoencephalography (MEG) studies have shown that the integration of visual and auditory or visual and tactile information can be associated with changes in phase coherence (Cooke et al., 2019; Doesburg et al., 2008; Hipp et al., 2011; Wang et al., 2019). Furthermore, studies in humans have provided evidence for changes of crossmodal coupling during audiovisual speech perception (Keil et al., 2012; Lange et al., 2013) and during evaluation of the duration of audiovisual stimuli (van Driel et al., 2014). Evidence is available for the modulation of dynamic crossmodal coupling of brain signals by factors such as attention, memory, emotion, or expectation (Senkowski et al., 2014; Senkowski et al., 2008; Talsma et al., 2010). In the present study, we have addressed the modulation of such neural dynamics in the context of predictions on upcoming multisensory stimuli.

The role of neural oscillatory activity in predictive processing has been postulated early on and is supported by studies on unimodal processing (for review, see Arnal & Giraud, 2012; Bastos et al., 2012; Engel & Fries, 2010; Engel et al., 2001; Hyafil et al., 2015). It has been suggested that predictions regarding stimulus identity or stimulus timing may involve oscillatory mechanisms in different frequency ranges. While the latter may be associated with low frequency (delta, theta) oscillations, the former might predominantly involve oscillations in higher frequency (beta, gamma) bands (Arnal & Giraud, 2012; Engel & Fries, 2010). The relation between oscillatory activity and predictions has been addressed by only a few studies in the context of multisensory processing, for review, see (Arnal & Giraud, 2012; Jessen & Kotz, 2013). While some of these studies have addressed temporal predictions (Arnal et al., 2015; van Wassenhove & Grzeczkowski, 2015), the dynamics underlying prediction of multisensory stimulus sequences has not yet been studied. Furthermore, large-scale dynamic coupling of oscillatory activity across brain areas has not yet been investigated in the context of multisensory predictions.

Here, we used MEG to investigate the neural mechanisms underlying the prediction of multisensory stimulus sequences. We designed an experiment in which participants judged whether the final item in a trial matched or violated the established sequence structure. This was contrasted with a condition where participants determined whether the final item was identical to either the immediately preceding item or the one before it. We analyzed both coupling and power differences in the MEG signals between conditions using a data-driven, multi-dimensional clustering approach that made no a priori assumptions about specific time windows, frequency bands, or brain regions of interest. This approach allowed us to investigate how intrinsically generated oscillatory signals are involved in predictions of stimulus sequences, and to what extent the coupling between different brain areas may relate to predictive processing.

## 2 Materials and Methods

### 2.1 Participants

29 healthy volunteers (17 females, age 26.3 ± 4.2 years) participated in this study. They gave written informed consent prior to commencing the experiment. All participants received 10 €/hour remuneration, and an additional 30 € bonus if they completed all sessions. All volunteers had a normal or corrected-to-normal vision and were right-handed and free of psychiatric or neurological disorders. The experiment was carried out in accordance with the guidelines and regulations, approved by the ethics committee of the Medical Association Hamburg.

### 2.2 Experimental procedure

A green fixation cross (0.25° visual angle) was shown at the center of the screen, and participants were asked to maintain fixation during the stimulation. A set of 5-item-long binary visual-auditory sequences (32 possibilities) was presented to the participants (Fig. 1). Each item appeared for 150 ms, followed by a 550 ms break before the next item, both with the fixation cross. An item was a vertical or horizontal Gabor patch (10° visual angle, 0.5 cycles per degree), accompanied by a simultaneously presented sine wave tone which was either higher (2000 Hz) or lower (1800 Hz) in pitch. The volume of the tone was adjusted to 30 dB above the individual hearing threshold, ensuring sufficient salience without causing discomfort. In each trial, only one of the sequences would be presented and repeated; all sequences were presented with equal probability for all participants. The length of the trials was randomized between 8 and 20 items, with an approximately fixed hazard rate — the probability that the sequence would end after each item was roughly constant, making the endpoint unpredictable. In the prediction task, participants had to monitor all items and to indicate, by button press, whether the last item of the trial violated the sequence. The next trial would start 1.0 – 2.0 (mean 1.5) seconds after the button press. For different blocks of trials, participants had to attend to either the auditory or the visual stimuli while ignoring the respective other modality. Although only the stimulus changes in the attended modality defined the task-relevant sequence, the combination of visual and auditory features was kept constant except for the last item (Fig. 1). The probability of a sequence violation was 50% in either modality, and violations occurred independently across the visual and auditory modalities. In the control memory task, the physical stimuli were exactly the same as in the prediction task, but participants had to indicate whether the last item was the same as the item second (1-back) or third (2-back) from last in either the visual or the auditory modality. The inclusion of both the easier 1-back and more difficult 2-back conditions aimed to minimize potential confounding effects of task difficulty between the prediction and memory tasks. For the formal analysis, the two conditions were combined, as subsequent analysis revealed no substantial differences in the MEG data between them. Overall, the study had a two by two design: task (prediction or P, vs. memory, or M) x relevant modality (visual or V, vs. auditory or A). There were 32 test trials in a block, and each participant accomplished 16 blocks for one recording session. Each block was initiated by the participants, who usually took a 0.5 ∼ 2 minutes break between blocks. A mandatory break of about 20 minutes was taken in the middle of each session. The order of blocks in a session was counterbalanced with respect to task and modality across participants. For each participant, two sessions were recorded on separate days, no more than 6 days apart. Before the first MEG recording session, a practice session comprising 4 blocks was conducted outside of the MEG chamber to familiarize participants with the task. The combination of visual and auditory features was different between the practice session and the MEG recordings. Only the data from the MEG sessions were used for subsequent analysis.

**Fig. 1.**
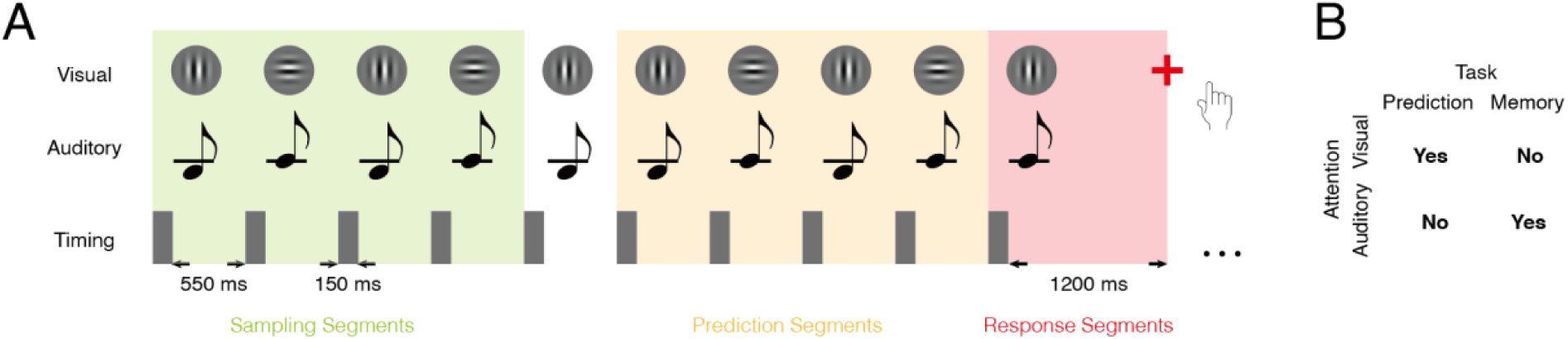
Experimental paradigm. (A) Trial structure: visual gratings (upper row) and auditory tones (middle row) were presented simultaneously in a sequential manner (timing in bottom row). A green fixation point was always presented in the center of the gratings during the presentation. The first five items constituted the sample sequence, which was repeated in the trial, with the exception of a potential change in the last item. Note that the last repetition could comprise less than five items: each trial comprised a total of 8 to 20 items. The number of items was randomized across trials, while hazard rates remained constant. When the fixation cross changed its color to red, subjects had to respond by button press. Data were segmented according to the onset of each stimulus and grouped into three categories: sampling segments (1^st^ to 4^th^ items), prediction segments (6^th^ to penultimate item), and response segments (final item). Each stimulus lasted 150 ms, followed by a 550 ms interval. The fixation cross turned red, as a response cue, 1200 ms after the last stimulus. (B) Correct answers for the example trial shown in (A). In the prediction task, subjects had to judge whether the last item matched with the sample sequence in the attended modality; in the memory task, subjects had to indicate whether the last item was identical to the item before (N-back) in the attended modality. In each block, the subjects performed one of the two tasks and attended to either the visual or the auditory modality. The order of tasks and attended modalities was counterbalanced across subjects.

The main experiment was conducted in a dimly lit, sound attenuated MEG chamber. The participants were seated in a recording chair with armrest. Visual and auditory stimuli were generated from the same computer (Dell Precision T5500) equipped with Psychtoolbox-3 (Brainard, 1997) and Matlab (R2017b, MathWorks, Natick, MA), on a Windows 7 Professional 64-bit operating system. The visual stimuli were delivered from a projector (VPX-PRO-5050B, VPixx Technologies Inc, Saint-Bruno, Canada) outside of the chamber via a mirror system to a thin film screen mounted 60 cm in front of the participants’ eyes, with a 100 Hz refresh rate. The auditory stimuli were delivered with MEG-compatible in-ear headphones (SRM-2525, STAX Limited, Fujimi, Japan). To ensure precise synchronization of auditory and visual stimuli, we used a custom-built a battery-powered device. A microphone was placed within the ear tube to capture the auditory signal; and a photodiode was positioned at the center of the screen to detect the onset of the visual stimulus. Both signals were converted to voltage and fed into auxiliary EEG channels integrated with the MEG acquisition system. We recorded these signals before the experiment and analyzed them offline to assess the delay between modalities and the temporal jitter across trials. Any systematic delay due to hardware latency was compensated in the stimulation code to ensure that auditory and visual onsets were synchronized in physical time. The button presses were recorded and merged to MEG data with an fORP system (HHSC-2X2, Current Designs Inc., Philadelphia, USA).

### 2.3 MEG recording and preprocessing

MEG signals were recorded with a 275-channel whole-head system (CTF MEG International Services LP, Coquitlam, Canada) in a magnetically shielded chamber, together with electrical eye, and cardiac activity via Ag/AgCl-electrodes in order to control for endogenous artifacts. Three magnetic sensors fixed at nasion, left and right preauricular points were used to track the head position, which was used for both online monitoring (Stolk et al., 2013) and offline artifact rejection. At the beginning of each block, participants were asked to reposition their head to the original position if the deviation was more than 10 mm. The recorded signals were digitized at a sampling rate of 1200 Hz. After removing malfunctioning channels, we obtained data from 274 sensors for each participant.

Raw data were read into Matlab with functions provided by FieldTrip version 20151002, (Oostenveld et al., 2011). Extreme noisy events (e.g., squid jumps, Oostenveld et al., 2011) were detected, and data within 1-second epochs around the event were excluded from further analysis. Data were then bandpass filtered between 0.5 ∼ 140 Hz with a Butterworth filter at the order of 4, in both forward and reverse direction for zero phase distortion. Z-score based automatic artifact detection (threshold z > 7, empirically determined from visual inspection of pilot data) was applied to label noisy segments for exclusion. The results were then visually inspected, with any remaining artifacts not captured by the algorithm manually marked. The average head position of each session of each participant was computed and used for later source reconstruction. Epochs where the head position deviated from this position more than 5 mm were also excluded. Instead of a notch filter, we removed 50 Hz line noise with sinusoidal signal estimation (Mitra & Pesaran, 1999). For artifact detection, ICA (Hyvarinen, 1999) was applied to the remaining data. To decide on the number of ICA components, a PCA was performed (Hyvarinen, 1999). We took the minimum number n, such that the first n PCA components preserved at least 99% of the whole information. Eye movements, cardiac artifacts, remaining 50 Hz line noise, and muscle activities were identified with their distinct spatial extension, spectral signature, waveform properties, and distribution across trials (Hipp & Siegel, 2013) and then removed from the data.

On average, 91.3 ± 0.8% of the original data were retained after artifact rejection. Following the exclusion of incorrect responses, and stratification across conditions, the number of EEG epochs used in the main coupling analysis was 715.7 ± 25.5 for the sampling segments and 767.7 ± 26.5 for the prediction segments.

### 2.4 Event extraction and MEG data segmentation

Time points of stimulus onsets and responses were extracted from the MEG data, and consistency with the events recorded by the stimulus software was verified. Behavioral performance such as hit ratio and response time was calculated based on these events.

The MEG data were segmented based on these events in several categories: (i) Onset segments, ranging from -700 to +700 ms relative to the beginning of each trial. This category was used to test neural responses to the physical stimulus properties compared to the pre-trial baseline. (ii) Sampling segments, defined as 700 ms epochs (150 ms stimulus + 550 ms inter-stimulus interval) time-locked to the onset of the 1^st^ to 4^th^ items of each trial. These segments captured the period during which participants acquired the sequence. (iii) Prediction segments, defined as 700 ms epochs after the onset of each item, from the 6^th^ to the penultimate item of each trial. This category was used to investigate how the participants predicted the sequences based on the acquired information. We did not include data segments around the 5^th^ item in the analyses because there might be confounding components for both learning and predicting: after the onset of the 5^th^ item, sampling information on this item and predicting the next one could not be disentangled. (iv) Response segments, ranging from 0 to 1500 ms relative to the onset of the last item. These segments were used to analyze response-related aspects. Any segment that overlapped with time intervals previously marked as artifacts was excluded from further analysis. Data were down-sampled to 400Hz after segmenting the clean data.

### 2.5 Behavioral analysis

Based on the events extracted from the MEG data, behavioral performance measures were computed, including hit rates, misses, correct rejections, and false alarms. Corrected accuracy was calculated as:

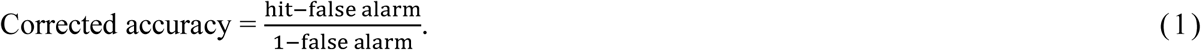

Response times (RTs) were also analyzed, considering only correct responses. RTs exceeding three standard deviations from the mean were excluded to minimize the influence of outliers.

A repeated-measures analysis of variance (ANOVA) was conducted on both corrected accuracy and response time using SPSS software. The ANOVA included two within-subject factors — task (prediction vs. control) and attended modality (visual vs. auditory) — and one between-subject factor— control condition (1-back vs. 2-back). Greenhouse-Geisser correction was used when necessary.

### 2.6 MRI data acquisition and pre-processing

T1-weighted structural MRI scans were obtained for 27 of the 29 participants. MRI recording was performed with a Magnetization Prepared Rapid Gradient Echo (MP RAGE) sequence on a Siemens 3 T scanner (Magnetom Trio, Siemens, Erlangen, Germany). with the following parameters: repetition time (TR) = 3150 ms, echo time (TE) = 1.37 ms, flip angle = 8°, bandwidth = 540 Hz/pixel, field of view = 260 × 260 mm, voxel size = 1.6 mm isotropic, and acquisition matrix = 160 × 160. The total acquisition time was approximately 13 minutes. The acquired images were converted from DICOM to NIfTI (Neuroimaging Informatics Technology Initiative) format and then processed using the FreeSurfer software package (Dale et al., 1999) for cortical segmentation, brain surface inflation and head shape reconstruction. Subsequently, the nasion and the left and right preauricular points were manually labeled on the reconstructed head shape and co-registered with coordinates of these three landmarks obtained from the MEG system with the aid of the MNE software package version 2.74 (Gramfort et al., 2014). This procedure put the MEG sensors and the structural brain into the same coordinate system.

### 2.7 Source reconstruction

We constructed our source space based on the FreeSurfer averaged template brain (Dale et al., 1999). To generate the source space, we started from an icosahedron and then recursively subdivided its edges, the middle point of which would be repositioned to its circumscribed sphere as new vertices. The defined sphere, with 162 vertices on it, was morphed onto the outer surface of the white matter (i.e., the inner surface of the gray matter) of each hemisphere, which yielded 324 locations homogeneously covering the entire cortex. By excluding locations that were unlikely to elicit significant brain responses (e.g., the corpus callosum), we reduced the number of source locations to 290 (Fig. S1). This procedure was accomplished with the MNE software package (Gramfort et al., 2014). We chose a relatively low number of source locations for reasons of computational efficiency and in order to be compatible with the physical spatial resolution of the MEG sensors, which varies by location but is typically on the order of several centimeters (Hauk et al., 2022). These source locations were then mapped to each individual anatomical MRI. For those participants without an anatomical MRI, we used the template brain ICBM152 from the Montreal Neurological Institute (Mazziotta et al., 1995) instead. As a next step, we computed lead fields with these source locations and individual sensor positions for each session. Lead field calculation was realized with a single-shell head model (Nolte, 2003). We did not apply compensation for ICA component reduction with the assumption that the lead field distribution was not altered by removing components related to eye movements or other muscle activity (Hipp & Siegel, 2015). Making use of linearly constrained minimum variance (LCMV) methods (Van Veen et al., 1997), we projected the original sensor-level waveforms to the source locations defined above. For each source location, we computed three orthogonal filters (one for each spatial dimension) that passed activity from the location of interest with unit gain while maximally suppressing activity from all other locations. Next, we linearly combined the three filters into a single filter in the direction of maximal variance (Hipp et al., 2011). In order to avoid spurious effects resulting from unequal filters in between-condition comparisons, we used data from all conditions (with the same number of trials for each condition) to compute a common filter. To derive source estimates, we multiplied the cleaned sensor data with the real-valued filter. Then we merged data from different sessions of the same participant. High source correlations can reduce source amplitudes estimated with beamforming due to source cancellation (Hipp et al., 2011; Van Veen et al., 1997) which might influence the magnitude of cortico-cortical coupling estimates. However, within the range of physiological source correlations (Leopold et al., 2003), the identification of cortico-cortical coupling using the beamforming method is possible (Kujala et al., 2008). Moreover, as source cancellation might reduce the sensitivity to detect coupling, it should not lead to false positive results.

### 2.8 Statistics

In general, we performed ANOVA for statistics on multiple factors, and paired t-test for pairwise comparison within one factor, in the analyses of following MEG data. Besides providing significance measures (*p* or *F* values), we also show measures of effect sizes (*η^2^* for ANOVA or Cohen’s *d* for pairwise comparison) where appropriate.

### 2.9 ERF computation

For each task (prediction, memory) and modality (visual, auditory) combination (i.e., four conditions), we averaged the waveforms in source space, with trial number stratified for the conditions, to obtain event-related fields (ERF) during each type of segment (onset, sampling, prediction, response).

### 2.10 MEG time-frequency decomposition

The time-frequency decomposition was performed in cortical source space. We computed an averaged ERF of all conditions in the onset segments, from which we evaluated the evoked activity by physical stimuli. All spectral estimates were obtained with the multi-taper method (Mitra & Pesaran, 1999) and computed across 19 logarithmically scaled frequencies from 5.66 to 128 Hz with 0.25-octave steps. We removed the ERF of respective conditions, from each data segment prior to further spectral analysis of the induced (non-phase-locked) activity. In the induced data, we took time points in steps of 100 ms (the time range of each segment was defined as described in “Event extraction and MEG data segmentation”). In the evoked data, we took time points in steps of 25 ms. We characterized responses relative to the pre-stimulus baseline using the bin centered at -100 ms. The temporal and spectral smoothing was performed as follows. For frequencies larger or equal to 16 Hz, we used temporal windows of 250 ms and adjusted the number of Slepian tapers to approximate a spectral smoothing of 3/4 octave; for frequencies lower than 16 Hz, we adjusted the time window to yield a frequency smoothing of 3/4 octaves with a single taper (Hipp et al., 2011). In case the window extended outside of the segment range, zero padding was applied. The employed time-frequency transformation ensured a homogenous sampling and smoothing in time and frequency, as required for subsequent network identification within this multi-dimensional space.

### 2.11 Power estimation and phase coupling analysis

Power of oscillatory signals was estimated from the spectrum *X (f, t)* as follows:

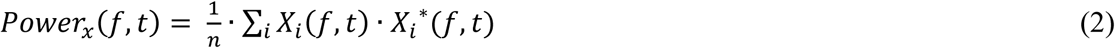

Here *i* is the trial index, *t* is the time bin index, and *f* is the frequency bin index, * represents the complex conjugate.

To quantify frequency-dependent synchronization in source space, we estimated the imaginary part of coherency (*ImC*) – a measure with minimum influence of volume conduction (Nolte et al., 2004). The *ImC* between pairs of signals *X (f, t)* and *Y (f, t)* was computed according to the following equation:

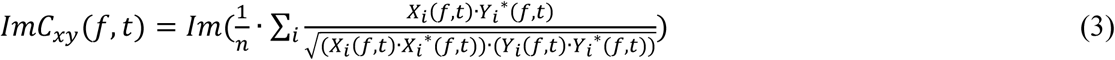

Here *i* is the trial index, *t* is the time bin index, and *f* is the frequency bin index, * represents the complex conjugate, *Im* indicates imaginary part of complex numbers.

### 2.12 Identification of networks by clustering

The general approach of our network identification closely resembles the method described in (Hipp et al., 2011; Wang et al., 2019) and is summarized below. An interaction between two cortical areas can be represented as a point in two-dimensional space and can be extended to four-dimensional space by adding the dimensions time and frequency. Identifying networks of significant interaction is then equivalent to identifying continuous clusters within this 4-dimensional space. In our approach, we first computed the ImC for all pairs of locations in all time and frequency bins and conditions with equation (3). Then t-statistics (for pair-wise comparison) or ANOVA (for multiple-factor analysis) was performed across participants. The resulting multi-dimensional matrix was then thresholded with a *t* or *F* value, setting every connection to either 1 (connected) or 0 (not connected). Further, we performed a neighborhood filtering using threshold to remove spurious connections: if the existing neighboring connections of a certain connection were fewer than certain ratio of all its possible neighboring connections, this connection would be removed. The specific threshold values used for each analysis are reported in the corresponding sections below. Each functional connection was defined in four dimensions: frequency, time, and the two source locations forming a pair. Two connections were considered as neighbors if they were identical in three of the four dimensions, while differing by only one step in the fourth—i.e., 100 ms in time, 0.25 octaves in frequency, or immediate neighbor in source space. Two source locations were defined as immediate neighbors if they were directly connected in the regular polyhedron during source space construction. Eventually, the remaining neighbor connections formed the candidate clusters for statistics. We defined their sizes as the integral of the *t* or *F* values across the volume, time and frequency. To evaluate their significance, cluster-based permutation tests were applied to account for multiple comparisons across the interaction space (Maris and Oostenveld, 2007). To this end, the cluster-identification approach was repeated 1000 times with shuffled condition labels to create an empirical distribution of cluster sizes under the null-hypothesis of no difference between conditions (Nichols & Holmes, 2002). The null-distribution was constructed from the largest clusters (two-tailed) of each resample, and only clusters with sizes ranked top 5% in these null distributions were considered as significant (i.e., *p* = 0.05) in the comparison. The final clusters corresponded to cortical regions with different synchronization status among comparisons, and were continuous across time, frequency, and pairwise space.

The identification of power difference clusters was similar to the above except that there was only one dimension in space rather than two for coupling.

Using the methods described above, we applied the cluster identification approach for the analysis of several contrasts. We identified the cluster of regions showing stimulus-related power changes by contrasting the evoked power from each time bin between 0 and 700 ms after the onset of the first item of each trial, with the power averaged between time bins between 125 and 75 ms before stimulus onset, with a pairwise t-test. Here, we use a threshold of *t* values corresponding to *p* = 0.01; the threshold for the neighborhood spatial filter was 0.3, i.e., at least 30% of neighboring difference had to be significant.

To investigate differences of induced oscillation power between conditions, we employed a 2 x 2 design ANOVA, with factors task (prediction vs. memory) and attended modality (visual vs. auditory). Here we use a threshold of *F* values corresponding to *p* = 0.05 for the main effects and interactions. The threshold for the neighborhood spatial filter was 0.3. This means that a test with a significant difference would be still regarded as significant after filtering, only when at least 30% of the neighboring tests were significant. *T*-test was applied for each component of the cluster to test the direction of the difference. If directions of effects were not consistent within a cluster, the positive and negative effects were analyzed separately. The same approach was used to investigate differences of coupling between conditions.

In order to investigate the effects of stimulus context, we classified the segments according to its previous or next stimulus, which could be same or different. With this factor and the attended modality, we construct another 2 x 2 ANOVA design. Cluster-based permutation test based on *F*-statistics (with the threshold of *F* values corresponding to *p* = 0.01) was applied for the main effects and interactions; the threshold for the neighborhood spatial filter was set to 0.3, i.e., at least 30% of neighboring differences had to be significant. Again, a post-hoc test was applied for each component of the network to test the direction of the difference. The positive and negative components within the clusters derived from the *F*-statistics were analyzed separately.

Furthermore, we tested both the power and coupling difference between sequence continuation and violation using paired *t*-tests. Cluster-based permutation test based on these results (with the threshold of *t* values corresponding to *p* = 0.01) was applied. In this case, the threshold of neighborhood spatial filter was also set to 0.3.

### 2.13 Illustration of identified clusters

To visualize networks identified by the clustering approach, we projected them separately onto two subspaces. We integrated the *t* or *F* values among connections over all spatial locations for each time and frequency bin to illustrate, when and at which frequencies a cluster was active irrespective of the spatial distribution. Complementary to this, we also computed the integral of *t* values in the connection space over time, frequency, and target locations (only for coupling clusters) for each location in the corresponding cluster. The result was then displayed on the fsaverage template brain surface to reveal the spatial extent of the cluster independent of its intrinsic coupling structure and time-frequency characteristics. For illustration purposes, the data were interpolated from 290 to ∼300,000 vertices on the inflated brain surface. The labeling of brain areas involved in our clusters was based on their relative position to the gyri and sulci of the template brain (Fischl et al., 2004).

### 2.14 Performance modulation by theta phase within the prediction network

We tested the modulation of behavioral performance by the phase of the coupling in the theta band clusters related to prediction (Fig. 5C and 5D). We defined the optimal phase relation of the coupling by averaging the segments used to quantify the cluster (all corrected and stratified for conditions) for each component. Each segment was classified into one of the eight bins based on its delay to the optimal phase relation. We averaged the response time from the same data for each component and then averaged these across the whole cluster. Since here the delay was a relative measure (difference to optimal), we averaged the tuning of response time by phase across participants.

We performed a similar analysis as the above to test for effects on accuracy, with two main differences as follows. First, we had to include all the data for computation of the percentage of hits, not only the data that were used to generate the network (corrected and stratified). Second, we used half of the data with correct responses, to generate the optimal phase relation and classify half of the full data, which would not include the trials to generate the optimal phase relation, and this was done as well for the respective other half of the data.

To rule out an influence of the number of segments, which was significantly larger for zero phase delay than a delay of π, we re-classified the data by absolute values of phase delay to the optimal phase relation, where the spacing was not exactly evenly distributed, e.g., the data range was smallest for the bin closest to zero so that it could include the same number of segments as the bin farthest from zero, which was much wider. Then a non-parametric correlation was computed to test whether the data followed the same trends. Spearman’s rho was calculated.

We also took the top and bottom 25% of data based on the absolute values of difference to optimal phase relation, and then compared the difference of performance with a paired t-test. We pooled the *t* values for each voxel. Then we also took the *F* values of each voxel when generating the cluster. The two groups of values were correlated, and Pearson’s *r* was measured.

### 2.15 Cluster analysis based on stimulus context

As mentioned in section “Identification of clusters for various contrasts”, we detected two clusters related to stimulus context, i.e., the occurrence of the same versus different stimulus as the previous or next item respectively. We calculated the modulation index (MI) by dividing the difference of ‘same’ and ‘different’ conditions and then dividing by their sum. MIs were also computed for the corresponding data for sampling segments and memory task, and compared using a 2 x 2 ANOVA.

### 2.16 Control condition analysis

For the identified clusters, we examined the interactions between control conditions (1-back and 2-back) and the primary contrast of interest, i.e., prediction vs. memory, in an ANOVA to rule out potential confounding effects of task difficulty.

### 2.17 Relation between clusters and cross-frequency coupling

To investigate interactions among clusters, especially between the theta coupling cluster and the clusters defined by the stimulus context, we computed their cross-frequency coupling. We first chose the locations that were most significant in both clusters, i.e., at least 1 standard deviation higher than the mean for the integrated *F* values. Then we computed the theta (5 - 7 Hz) phase of these voxels in the theta networks and the beta (12 - 32 Hz) power of these voxels in the beta clusters. The cross-frequency coupling among these time courses was computed with the modulation index (Tort et al., 2010; Wang et al., 2019). As a control, we computed a shift predictor by shuffling the trials, such that the phase from trial A would be coupled with trial B, with all other intact. In this case, the stimulus-locked information would be preserved but the instantaneous correspondence of phase and amplitude was removed. Subsequently, the cross-frequency coupling of real data was normalized with those obtained from shift predictors. Then they were compared across conditions with ANOVA.

## 3 Results

### 3.1 Behavioral data

Participants performed the task with an average corrected accuracy of 90.9%. No significant three-way interaction (*F* (1, 27) = 0.04, *p* = 0.85, *η^2^p*< 0.01) among the factors (task, attended modality, and control condition) was observed with regard to corrected accuracy. Additionally, there were no significant two-way interactions between any two factors (task and attended modality: *F* (1, 27) = 1.49, *p* = 0.23, *η^2^p*= 0.05; task and control condition: *F* (1, 27) = 2.91, *p* = 0.10, *η^2^p*= 0.10; attended modality and control condition : *F* (1, 27) = 3.01, *p* = 0.09, *η^2^p*= 0.10). The main effect of attended modality was also not significant (*F* (1, 27) = 0.14, *p* = 0.71, *η^2^p*< 0.01). However, the main effect of task was significant (*F* (1, 27) = 4.49, *p* = 0.04, *η^2^p*= 0.14), with corrected accuracy being higher in the memory task compared to the prediction task (Fig. 2). A significant between-subject effect of control condition was also observed (*F* (1, 27) = 3745.56, *p* < 0.001, *η^2^p*= 0.99), with higher corrected accuracy for the 1-back control compared to the 2-back control group.

**Fig. 2.**
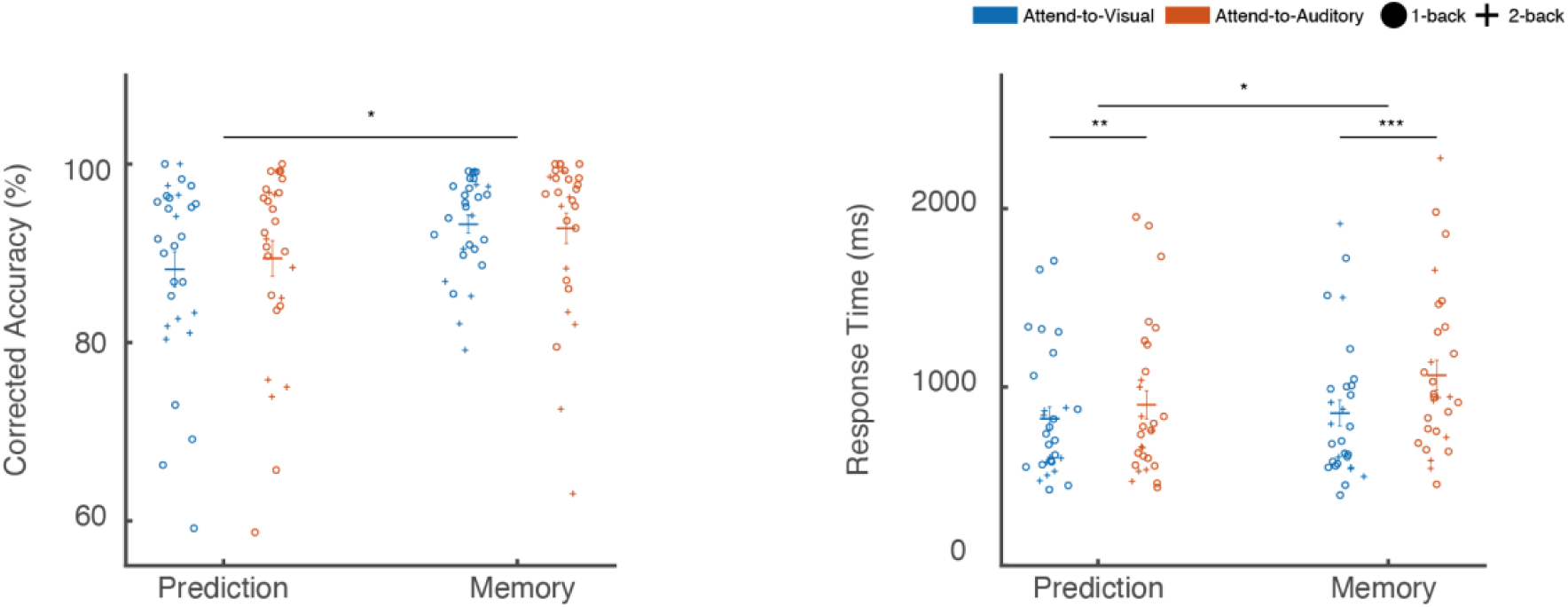
Behavioral performance. The left panel shows corrected accuracy (%) and the right panel shows response times (ms) for the prediction and memory tasks. Results are further classified by attended modality (Attend-to-Visual *vs.* Attend-to-Auditory, color coded) and control condition (1-back *vs.* 2-back, symbol coded). Horizontal bars represent group means, and vertical bars indicate standard errors of the mean (*n* = 29). Each circle or cross corresponds to an individual participant’s performance. Significant differences are indicated: * *p* ≤ 0.05, ** *p* ≤ 0.01, *** *p* ≤ 0.001

For response times, no significant three-way interaction was found (*F* (1, 27) = 0.27, *p* = 0.60, *η^2^p*= 0.01), nor was there a significant interaction between control condition and attended modality (*F* (1, 27) = 0.58, *p* = 0.45, *η^2^p*= 0.02). However, a significant interaction between task and attended modality was observed (*F* (1, 27) = 9.55, *p* < 0.01, *η^2^p* = 0.26). Post-hoc analysis revealed that in the prediction task, participants responded significantly slower when attending to the auditory modality (*p* < 0.01, Cohen’s *d* = 0.20). In the memory task, this effect was even larger (*p* < 0.001, Cohen’s *d* = 0.50) (Fig. 2). A significant interaction between task and control condition was also observed (*F* (1, 27) = 11.77, *p* < 0.01, *η^2^p*= 0.30). Post-hoc analysis showed no significant difference in response times between tasks in the 1-back control group (*F* (1, 27) = 0.02, *p* = 0.90, *η^2^p*< 0.01). However, in the 2-back control group, responses were significantly faster in the prediction task compared to the memory task (*F* (1, 27) = 7.01, *p* = 0.03, *η^2^p* = 0.47). Finally, a significant between-subject effect of control condition was found (*F* (1, 27) = 135.99, *p* < 0.001, *η^2^p*= 0.83), with faster responses observed for the 1-back control group.

### 3.2 Audiovisual stimuli lead to power changes in multiple frequency ranges

Since in all conditions we always presented similar audiovisual stimuli (gratings accompanied by tones), we pooled the data across conditions with stratified trial numbers and examined the power change caused by the first stimuli of the sequences, as compared to baseline (100 ms before trial onset) with the clustering approach. The results are illustrated in Fig. 3. The stimuli generally caused a power increase in most of the brain areas, which was most pronounced in bilateral occipital areas, fusiform gyri, transverse temporal gyri, superior temporal gyri, superior temporal sulci, intraparietal sulci, precunei, and cingulate cortices. The stimulus-related activation occurred in a broad range of frequencies. The activations were strongest after stimulus onset (0 ms) and showed a second peak after stimulus offset (150 ms). The spatial and temporal distribution of these effects is consistent with typical early sensory responses and supports the validity of our source reconstruction.

**Fig. 3.**
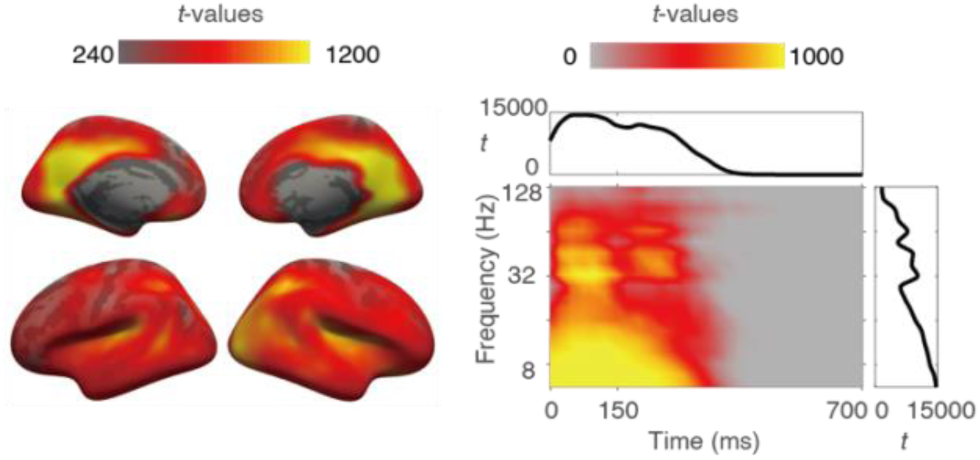
Stimulus-related power changes. (Left) Spatial distribution of the cluster, see method for details. Color intensity is scaled to the integral of *t* values across time bins and frequency bins for each voxel. (Right) Time-frequency distribution of the cluster; color intensity is scaled to the integral of *t* values across all voxels for each time and frequency bin. Time 0 indicates stimulus onset. Inset plots show integrated *t* values over time (top, x-axis: time in ms, y-axis: *t* value) or frequency (right, x-axis: *t* value, y-axis: frequency in Hz).

### 3.3 Prediction task induces less beta power than memory task

We observed main effects regarding power differences between prediction and memory tasks in both sampling (permutation test, *p* = 0.018, Fig. 4A) and prediction segments (permutation test, *p* = 0.015, Fig. 4B), in a 2 x 2 ANOVA design with the two factors task and attended modality. Since the interaction between these two factors did not yield any significant clusters, we consider the main effects separately. In this paper, we focus on the prediction-related dynamics and will present a detailed analysis of attention-related effects in a separate publication. The cluster showing a significant main effect of task on power differences involved bilateral superior temporal gyri, fusiform gyri, precentral gyri, insula, cingulate cortices, lateral orbitofrontal cortices, and pars opercularis. To determine the direction of the effect, we examined the average power within the significant cluster and found that beta power was consistently higher in the memory task than in the prediction task. In the sampling segments, significant differences were observed in the alpha (8–11 Hz) and beta (19–32 Hz) bands. The alpha component, observed primarily in occipital areas during the sampling segments, was absent in the prediction segments.

**Fig. 4.**
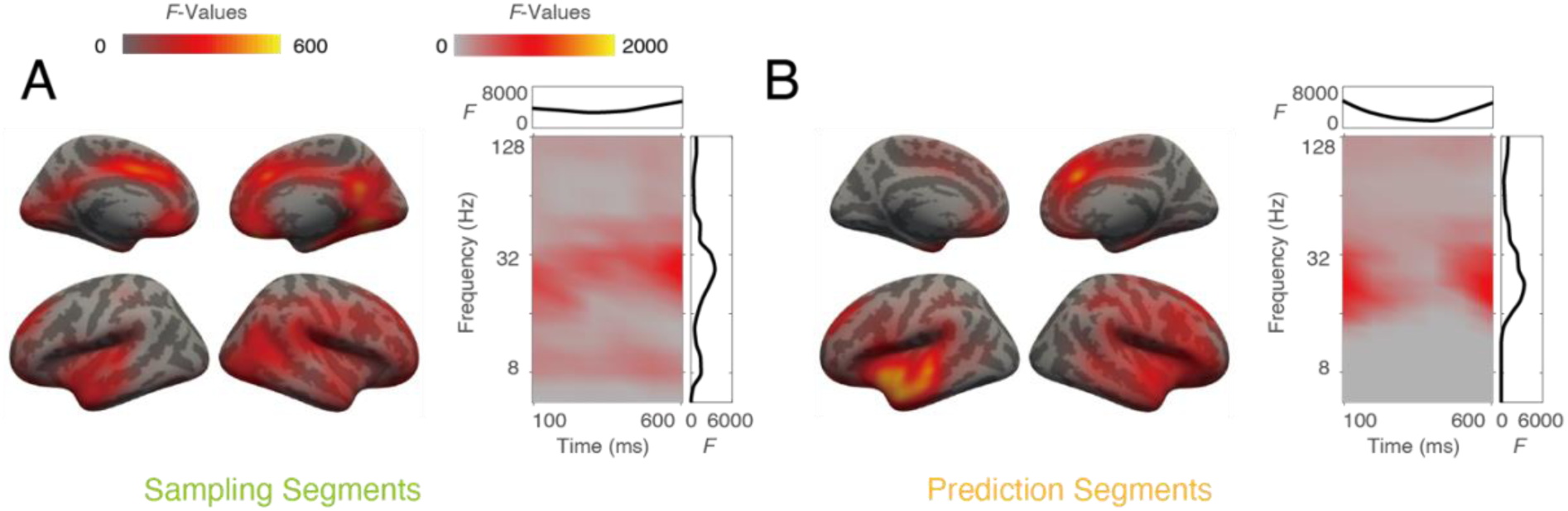
Power differences across tasks. The clusters were obtained with 2 x 2 ANOVA design on oscillation power, during both sampling (A) and prediction (B) segments. A significant main effect of factor task was observed. For each cluster, both spatial (left) and time-frequency (right) distributions are illustrated. Color intensity indicates accumulated *F* values. Side plots in the box are integrated *F* values plotted against time (top, x-axis: time in ms, y-axis: *F* value) or frequency (right, x-axis: *F* value, y-axis: frequency in Hz).

### 3.4 Prediction is associated with stronger theta coupling and weaker gamma coupling

As the next step, we analyzed functional connectivity, as measured by the imaginary part of coherency (Nolte et al., 2004), which minimizes the influence of volume conduction. We employed the same ANOVA design as for the power analysis in the preceding section. We observed significant coupling differences between prediction and memory task in both sampling (permutation test, *p* = 0.018, Fig. 5A) and prediction segments (permutation test, *p* = 0.026, Fig. 5B). The clusters were similar for both segment types and extended to superior temporal gyri, pre- and postcentral cortices, cingulate cortices, prefrontal areas, insulae, and precunei in both hemispheres. Coupling differences between the two tasks were observed in the theta (5 - 7 Hz), alpha (9 - 11 Hz), and gamma (> 70 Hz) frequency ranges. Interestingly, further analysis revealed that the theta and alpha coupling was stronger in the prediction task (Fig. 5C and 5D), whereas gamma coupling was stronger for the memory task (Fig. 5E and 5F). The spatial distribution of low-frequency components was similar to that of the high-frequency components. This suggests that, in these brain areas, theta and alpha coupling supports the prediction task, whereas gamma coupling may be involved in the memory task.

### 3.5 Control condition does not interact with task effects in identified clusters

For all these identified clusters, we examined the interactions between the between-subject factor control condition (1-back *vs.* 2-back) and the within-subject factor task (prediction *vs.* memory), which represented the primary contrast of interest. No significant interactions were observed in any of these analyses: beta power, *F* (1, 27) = 1.55, *p* = 0.22, *η^2^p* = 0.05; theta coupling, *F* (1, 27) = 1.98, *p* = 0.17, *η^2^p* = 0.07; gamma coupling, *F* (1, 27) = 0.01, *p* = 0.92, *η^2^p* < 0.001.

To visualize these results, Fig. S2 illustrates the differences between prediction and memory tasks for both power and coupling measures in respective clusters, with data further classified by control condition.

### 3.6 Cross-regional theta phase delay modulates behavior

It has been suggested that brain areas communicate via coherent oscillations when the interaction occurs at an optimal phase relation (Rohenkohl et al., 2018; Womelsdorf et al., 2007). Evidence supporting this hypothesis has been obtained from neurophysiological recordings in animal studies, specifically at the level of single neurons or small neuronal populations. In the present study, we tested whether this notion is also supported by the analysis of large-scale data recorded with MEG from the human brain. In the observed theta clusters (Fig. 5C and 5D), we determined the optimal phase relation for each component (as defined by a certain location pair, as well as time and frequency bin) as the average phase lag between the two locations at this time and frequency bin, in all segments corresponding to the data used to generate them (correct trials stratified for each condition). We then classified the phase delays relative to the optimal phase relation into 8 bins (from -π to π), where performance data (response time in the same dataset, and percentage of hits with all data included) were pooled and averaged. Thus, we could obtain the modulation of performance by phase delays for each cluster (Fig. 6A-6D). The results suggest that in both sampling and prediction segments, the percentage of hits was highest at zero delay to the optimal phase relation, and then decreased with increasing delay, forming a cosine-like function for the prediction task (Fig. 6A and 6C, red). Similarly, the response time of the prediction task was shortest at zero delay and then increased with phase delay for the prediction segments, resembling an inverted cosine function (Fig. 6D, red). The response time modulation in the prediction task during sampling segments followed the same trends but was less pronounced (Fig. 6B, red). The modulation of percentage of hit and response time in the memory task exhibited a similar but much weaker pattern in the prediction segments (Fig. 6C and 6D, green), but not in the sampling segments.

**Fig. 5.**
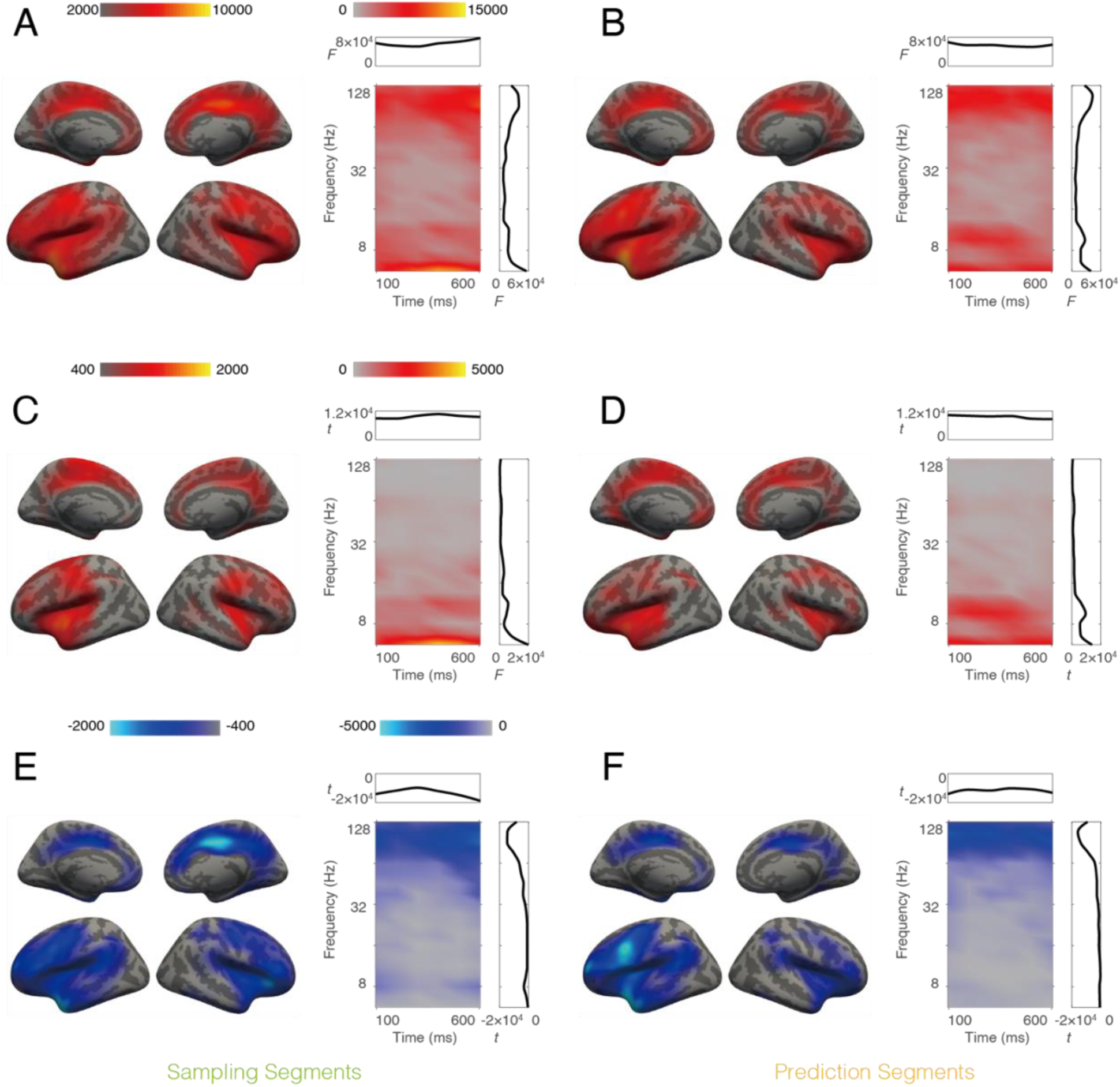
Coupling differences between prediction and memory tasks. (A, B) Main effects of the factor task in a 2 x 2 ANOVA design. (C-F) Results of paired t-test for the same contrast. The left column shows data from sampling segments and the right column shows data from prediction segments. In each panel, the spatial distribution of the cluster is displayed on the left. Color intensity is scaled to the integral of *F* or *t* values across time bins, frequency bins, and connected locations for each voxel. On the right, the time-frequency distribution of the cluster is shown, with color intensity scaled to integral of *F* or *t* values across voxel pairs for each time and frequency bin. Side plots in the box are integrated *F* or *t* values against time (top, x-axis: time in ms, y-axis: *F* or *t* value) or frequency (right, x-axis: *F* or *t* value, y-axis: frequency in Hz). Color bars are shared between panels A, B, C, D, and E, F.

**Fig. 6.**
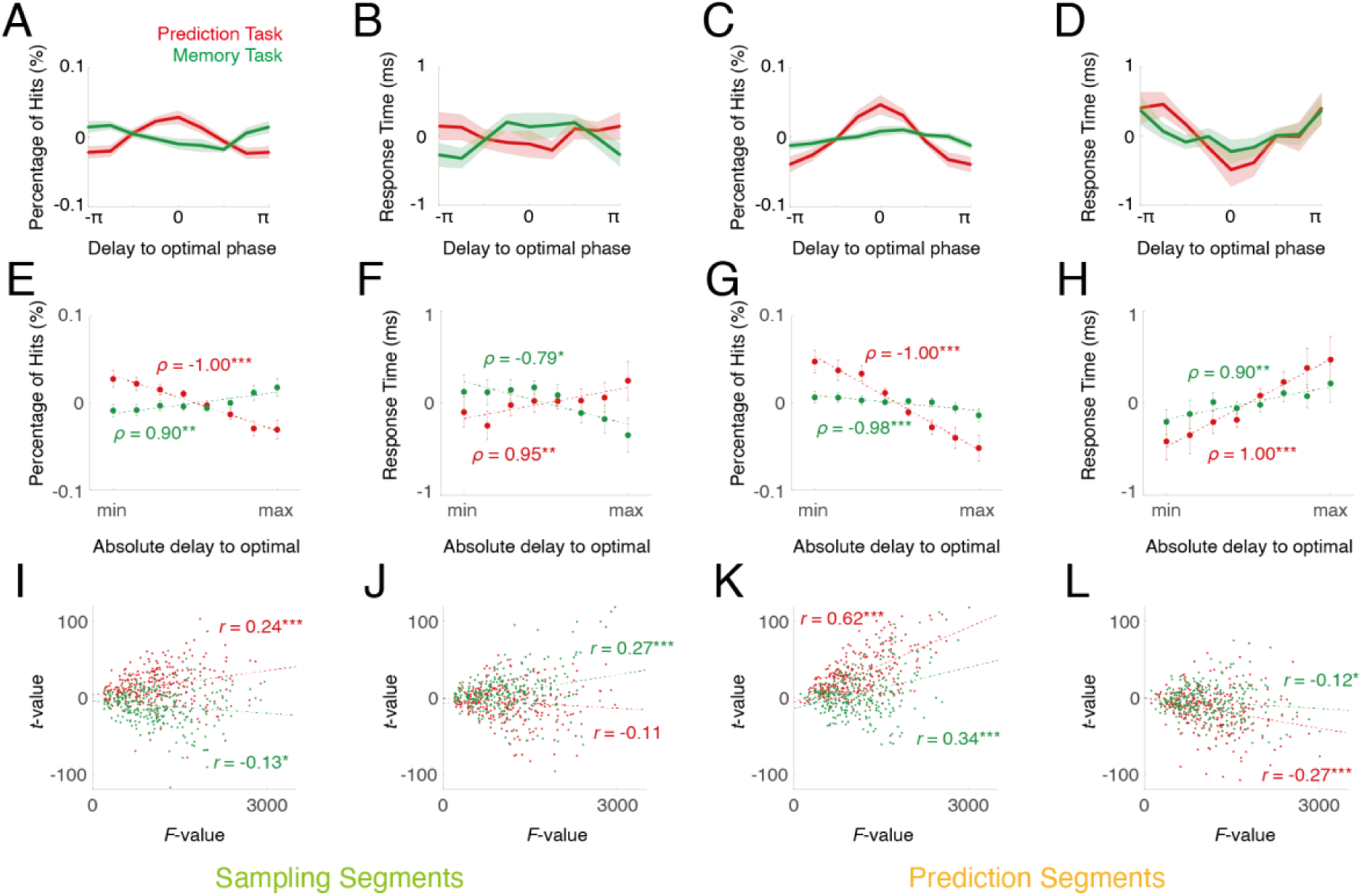
Dependence of behavior on theta coupling phase. Top row: dependence of percentage of hits (A, C) and response time (B, D) on phase delay to the optimal phase relation. Middle row: dependence of percentage of hits (E, G) and response time (F, H) on absolute delay to the optimal phase relation (n = 29), with an equal number of segments in each bin. Error bar indicates standard error. Spearman’s correlation coefficient (*ρ*) is indicated. In panel A – H the mean was removed for all percentages of hits and response times. Bottom row: Correlation between statistical power for percentage of hits (I, K) and response time (J, L) difference, and statistical power of clusters among the voxels. Pearson’s correlation coefficient (*r*) was indicated on the plots. The left two columns show data from the sampling segments while the right two columns show data from the prediction segments. \**p* ≤ 0.05, \*\**p* ≤ 0.01, \*\*\**p* ≤ 0.001

There were more data segments in the bin with zero delay to the optimal phase relation, and fewer in those with larger delays (e.g., close to -π and π). To rule out the possibility that the difference of phase modulation was due to the amount of data, we sorted the segments into 8 bins, according to absolute delay to the optimal phase relation, such that within each bin, there were equal number of segments. Subsequently, the performance per bin was tested with non-parametric correlation (Fig. 6E-6H), which confirmed that the effects were not caused by variations in the amount of data.

We further conducted a paired t-test on the data associated with the top 25% and bottom 25% delays for each voxel. These results were correlated with the integral *F* values of each voxel from the clustering approach, which yielded a significant positive correlation (Fig. 6I-6L). This analysis suggested that the more likely a brain location was part of the theta cluster, the stronger the behavioral modulation by the theta phase.

### 3.7 Beta power reflects stimulus memory and prediction

To successfully perform the prediction task, participants needed to remember the past stimulus and make assumptions about the next stimulus. To rule out the influence of the physical properties of the preceding or forthcoming stimulus itself, we analyzed a contrast defined by the stimulus context. Each segment was classified by its relation to the preceding segment, either the same or different, in the attended modality. This yielded a 2 x 2 ANOVA with factors modality (auditory vs. visual) and context (same vs. different). We applied our clustering approach with this design and identified one cluster for the main effect of context (relation to previous item, permutation test, *p* < 0.001, Fig. 7A). The cluster mainly involved lateral occipital areas, intraparietal areas, and precunei. It peaked around 400 ms in time and 13.5 Hz in frequency. Further analysis suggested that the power was stronger for the ‘same’ condition than the ‘different’ condition. We defined a modulation index (MI) as (same – different) / (same + different). The MIs were always larger than zero (all four comparisons p < 0.001, Cohen’s *d* = 1.41 for the prediction task in sampling segments, 1.32 for the memory task in sampling segments, 1.95 for the prediction task in prediction segments, 1.53 for the memory task in prediction segments, Fig. 7B), and were stronger during prediction segments than in sampling segments (*F* (1, 28) = 4.95, *p* = 0.034, *η^2^p*= 0.15, Fig. 7B). These results suggest that the information on preceding stimuli could be temporarily stored in this network.

**Fig. 7.**
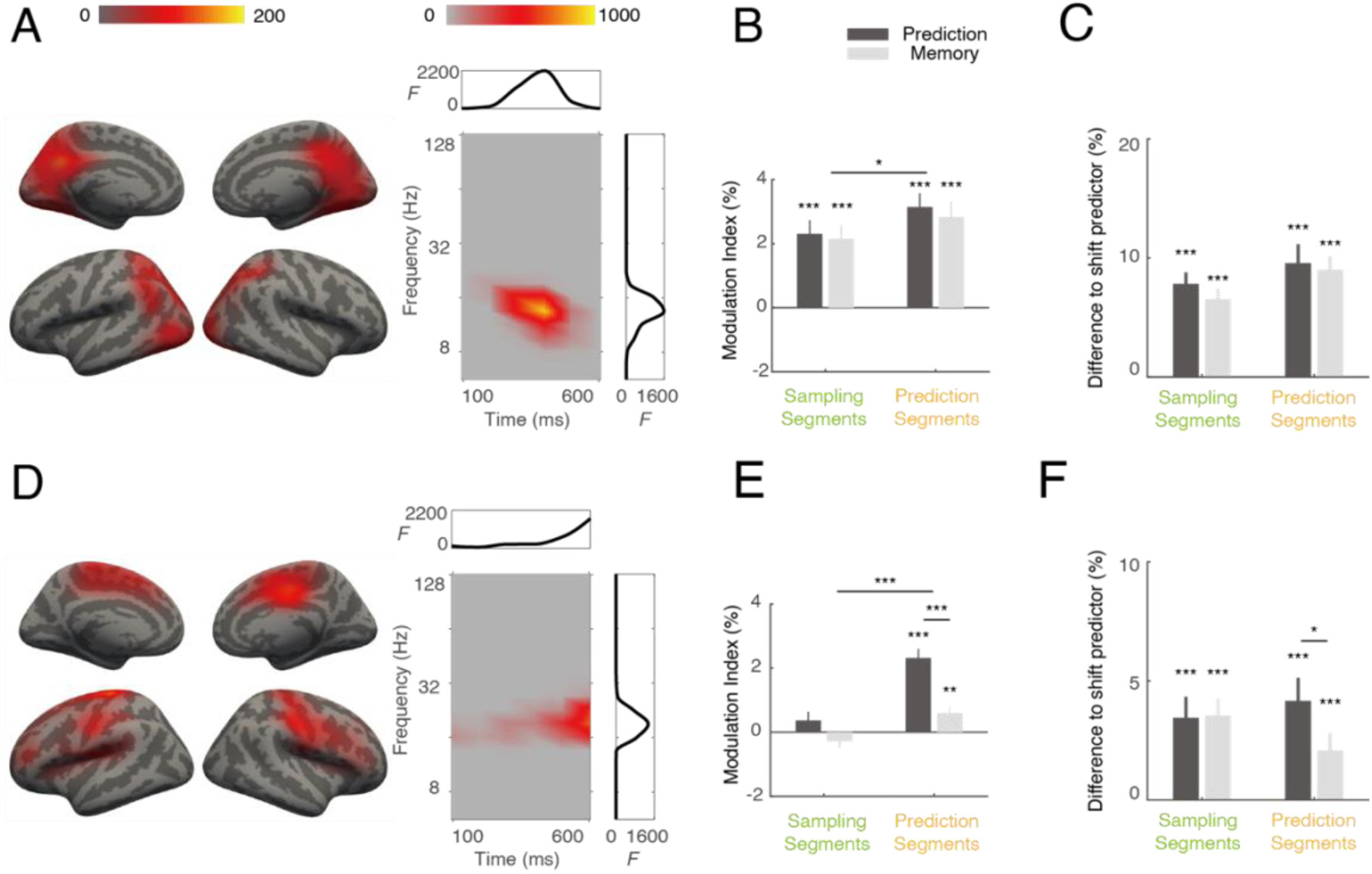
Power differences as a function of context. (A - C) Effects of same vs. different preceding item. (D - F) Effects of same vs. different subsequent item. Left column (A, D): spatial (left) and time-frequency (right) distributions of the identified cluster are illustrated. Color intensity indicates accumulated *F* values. Side plots in the box are integrated *F* values over time (top, x-axis: time in ms, y-axis: *F* value) or frequency (right, x-axis: *F* value, y-axis: frequency in Hz). Color bars are the same for panel A and D. Middle column (B, E): Modulation indices for same vs. different context conditions (n = 29). Right column (C, F): Phase amplitude coupling between significant locations shared between the coupling cluster and the context clusters, expressed as difference to the shift predictor (n = 29). \**p* ≤ 0.05, \*\**p* ≤ 0.01, \*\*\**p* ≤ 0.001

A similar analysis revealed another cluster for the main effects of context (relation to the next item; permutation test, *p* < 0.001; Fig. 7D). It mainly comprised pre- and postcentral areas, supplementary motor areas, and lateral prefrontal regions. It peaked around 600 ms after stimulus onset (or 100 ms before the next stimulus) at a frequency of 19.0 Hz. Further analysis suggested that the power was stronger for the ‘same’ condition than the ‘different’ condition. MIs in this cluster were not significantly different from zero in sampling segments but were significantly larger in prediction segments (*p* < 0.001, Cohen’s *d* = 2.17 for prediction task, *p* = 0.007, Cohen’s *d* = 0.76, for memory task, Fig. 7E). It was stronger for the prediction task than memory task (*F* (1, 28) = 27.01, *p* < 0.001, *η^2^p* = 0.49, Fig. 7E). This suggests that, before the onset of the next stimulus, activity in this network might encode the forthcoming sequence items.

### 3.8 Multiple clusters are linked by phase-amplitude coupling

A key question is how these different clusters relate to each other, i.e. how the respective neural populations interact to accomplish prediction. One possible mechanism is that the context from preceding items (prior knowledge) is accumulated to form a sequence representation, which is then used to predict the next item (consequence). Thus, the two beta power clusters may reflect the sources and outcomes of prediction, while the theta coupling cluster might serve as the link, mediating the predictive process. To test this idea, we examined cross-frequency coupling for the areas shared by these clusters, specifically investigating whether the theta phase of the coupling cluster modulated the beta power of the context cluster. We observed that in all cases cross-frequency coupling was stronger than a shift predictor computed as a control (Fig. 7C). Notably, the cross-frequency coupling was stronger for the prediction task than the memory task only in the prediction segments (Fig. 7F).

### 3.9 Sequence continuation vs. violation induces differences in both power and coupling

Finally, we examined the differences in power and coupling between sequence continuation and violation for the response segments using paired *t*-tests. The observed power differences are shown in Figure S3. The first cluster (permutation test, *p* < 0.001, Fig. S3A) exhibited stronger power for sequence continuation. The major frequency components of this cluster were in the theta-alpha range (5-12 Hz) and peaked around 900 ms. The activated areas in this cluster included the precunei, cingulate cortices, pre-and postcentral areas, caudal middle frontal areas, superior temporal gyri, middle temporal sulci, transverse temporal gyri, intraparietal sulci and insulae in both hemispheres, and superior and orbito-frontal areas in the right hemisphere.

The second cluster (permutation test, *p* < 0.001, Fig. S3B) involved the anterior cingulate cortices in both hemispheres, and superior temporal cortex, insular, pre-and postcentral areas, inferior precentral sulcus in the right hemisphere, where the theta (5-7 Hz) power was stronger for sequence violation after stimulus onset; this effect peaked around 300 ms.

The third cluster (permutation test, *p* < 0.001, Fig. S3C) included middle cingulate cortex, superior temporal cortex, insular, pre-and postcentral areas in the right hemisphere, where the beta (11-23 Hz) power was stronger for sequence violation just before the response cue; this difference was maximal around 1100 ms after stimulus onset.

We also examined the coupling difference, as measured by the imaginary part of coherency, between sequence continuation and violation for the responding segments with a paired *t*-test. The results are shown in Fig. S4. The first cluster (permutation test, *p* = 0.001, Fig. S4A) exhibited stronger coupling for stimulus continuation in the theta range (5-7 Hz), peaking around 1000 ms.

Activated areas in this cluster included middle cingulate cortices, pre-and postcentral areas and transverse temporal gyri in both hemispheres, and superior temporal gyri, insulae, and superior frontal gyri mainly in the right hemisphere.

The second cluster (permutation test, *p* = 0.005, Fig. S4B) included middle cingulate cortices, pre- and postcentral areas, inferior frontal areas and insulae in both hemispheres and superior temporal cortex mainly in the right hemisphere, where the theta (5-7 Hz) coupling was stronger for sequence violation after stimulus onset; this effect peaked around 400 ms.

## 4 Discussion

In the present study, we have recorded MEG during a serial prediction task in which participants had to acquire stimulus sequences and to monitor whether subsequent probe items complied with the sequence. Presentation of the audiovisual stimulus items revealed activation in a widely distributed set of areas involving early and higher-order visual and auditory areas as well as intraparietal sulci, precunei, and cingulate cortices. Task-related changes of power and coupling of neural oscillations were observed in several frequency bands. We identified prediction-related theta-band coupling in a network involving cingulate, premotor, prefrontal, and superior temporal regions (Figure 5). Behavioral performance of the participants correlated with phase delays between remote regions in this network (Figure 6). In this network, beta power was stronger when either the previous or upcoming stimuli matched the current item in the sequence. Furthermore, the amplitude of these beta signals was modulated by the phase of theta activity, as revealed by analysis of cross-frequency coupling (Figure 7). Overall, our findings suggest that oscillations in multiple frequency ranges, as well as coupling within and across frequency bands, play a crucial role in sequence processing. Given that no interactions between the prediction-related activity and attention-related activities were detected in our data, the attention effects observed with our paradigm will be presented in a separate paper.

Prediction is a key feature of brain function. Based on previous experience, the brain continuously generates predictions about upcoming events, which are then adapted through learning in order to reduce prediction errors (Bastos et al., 2012; Friston, 2010). A role of neural oscillations in predictive processing has been postulated early on and is supported by work on unimodal processing, for review see e.g. (Arnal & Giraud, 2012; Bastos et al., 2012; Engel & Fries, 2010; Engel et al., 2001; Hyafil et al., 2015). It has been suggested that predictions regarding stimulus identity or stimulus timing may involve oscillatory mechanisms in different frequency ranges. While the latter may be associated with delta and theta band oscillations, the former might be mediated by beta and gamma band interactions (Arnal & Giraud, 2012; Engel & Fries, 2010). It is important to note that the term *multisensory* can be interpreted in multiple ways. In the present study, we examined how predictive processing unfolds in a context where auditory and visual stimuli are always both presented. While this does not establish multisensory integration in the classical sense (e.g., behavioral enhancement over unisensory baselines), it does reflect prediction mechanisms under multisensory input conditions. In a separate, parallel study, we investigated a design in which successful task performance explicitly required integration across modalities, thus reflecting another aspect of multisensory processing.

So far, only relatively few studies have addressed the relation between oscillatory activity and predictions in the context of multisensory processing, for review, see (Arnal & Giraud, 2012; Jessen & Kotz, 2013). One of the earliest studies was carried out by Widmann and coworkers (Widmann et al., 2007), who used a matching paradigm in which visual score-like symbols predicted corresponding sound patterns. Compared to incongruent sounds, expected sounds evoked higher gamma-band activity. Van Ede et al (van Ede et al., 2010) presented subjects with a repeated series of concurrent tactile and auditory stimuli in which rare deviant stimuli occurred in one of the two modalities. Interestingly, prediction-related modulation of beta-band oscillations was observed even for unattended tactile stimuli. This modulation was enhanced during states of attentive expectation. Arnal et al (Arnal et al., 2011) investigated a violation of multisensory predictions in speech perception and demonstrated that correct predictions were associated with delta oscillations, whereas prediction errors were predominantly accompanied by beta- and gamma-band activity. Another recent study on crossmodal effects in speech perception (Biau et al., 2015) has observed modulation of theta-oscillations during auditory speech processing by concomitant visual input from observation of gestures. Low-frequency oscillations have also been found to be associated with temporal prediction in a study where a visual color change predicted the occurrence of an auditory stimulus (van Wassenhove & Grzeczkowski, 2015). A more recent study has observed a phase reset of delta oscillations in a task involving visual-to-tactile temporal predictions (Daume et al., 2021). Altogether, only a few studies are currently available that have addressed changes in oscillatory activity during multisensory prediction. Of course, the term multisensory may have various meanings. In this study we demonstrated how prediction was performed in a multisensory context. We have another parallel study where cross-modality information have to be integrated to perform the task, which reflect another aspect of multisensory.

To our knowledge, this study presents the first investigation of neural oscillations and their coupling in networks underlying prediction of multisensory stimulus sequences. In the analysis of task-related power changes, we observed significant differences between the prediction task and the memory control condition. The cluster reflecting these differences was mainly found in the beta range, comprising bilateral superior temporal gyri, sensorimotor cortices, insulae, cingulate and frontal cortices. In addition, there was an alpha component during sampling of the sequence but not in the prediction phase of the trials. This alpha component, which was mainly observed in visual areas, was more pronounced for the memory control task. This suggests that functional inhibition, as indicated by the alpha oscillations (Jensen & Mazaheri, 2010), was stronger during the sampling phase in the memory task. In this control condition, the information present in the sampling segments was task-irrelevant, whereas it was vital to construct the sequence in the prediction condition. Thus, it was not surprising that functional inhibition occurred for visual items in the memory task, while inhibition of visual processing was released during the sampling phase in the prediction task. Beta power was stronger during the memory task than during the prediction task, in line with the proposal that beta-band oscillations reflect the maintenance of the task-relevant information in a cognitive state(Engel & Fries, 2010). In the prediction task, the participants needed to constantly update the sequence information after the presentation of an item and predict the upcoming sensory inputs in the interval before the occurrence of the subsequent item; this cognitive updating might relate to the observed reduction of beta power.

In the same network, we also observed power differences in the responding segments, where subjects had to indicate whether the last item continued or violated the acquired sequence. These data suggest a beta power decrease upon violation of the sequence, with a subsequent beta rebound. This dynamic of beta power shows similarities to changes associated with prediction errors, as proposed in predictive coding frameworks (Arnal & Giraud, 2012; Bastos et al., 2012).

In addition to the task-related power differences, we also observed functional connectivity clusters defined by coupling differences between the prediction and the memory task. All clusters were similar in their spatial distribution, and they also shared a large proportion of the involved brain regions with the cluster obtained by comparing power difference. This suggests that these clusters, although formally derived in separate analysis steps, reflect prediction-related dynamics in the same cortical network. This prediction-related network comprised mainly superior temporal gyri, sensorimotor cortices, insulae, cingulate and frontal cortices. in both hemispheres. An interesting observation was that motor and pre-motor areas were quite prominent in the prediction network. This is in line with previous fMRI studies that have demonstrated a critical involvement of motor and premotor regions in prediction tasks (Arnal & Giraud, 2012; Bubic et al., 2010; Schubotz & von Cramon, 2002). In this network, coupling in the theta-alpha range was stronger for the prediction task compared to the memory task during both the sampling and the prediction phase of the trials, suggesting that coupling at low frequencies was essential for the prediction process. Interestingly, in the same network, coupling also occurred in the gamma frequency range, which was stronger for the memory condition. A possibility is that, in the memory task, the comparison of low-level visual or auditory features promoted gamma coupling, which has been related to feedforward transfer of sensory information(Bastos et al., 2015; Engel & Fries, 2010). In contrast, in the prediction task, the information on the 5-item sequence needed to be constantly retrieved and updated, which could require more feedback communication that may primarily be conveyed by coupling of low-frequency oscillations (Bastos et al., 2015; Fries, 2015). Interestingly, changes in theta-band coupling were also related to the processing of sequence continuation vs. violation in the response segments of the prediction trials.

We employed two types of control tasks—1-back and 2-back memory tasks—based on the assumption that the 1-back task would be easier than the prediction task, while the 2-back task might be overly difficult. By including both control conditions, we aimed to balance task difficulty and observed a combined outcome of more accurate but slower responses compared to the prediction task. While we hypothesized that the difficulty difference between the prediction and memory tasks would be minimal, this required confirmation from the data. The fact that the two control conditions showed significant differences in both corrected accuracy and response times indicates a clear distinction in difficulty. If the observed differences between the prediction and memory tasks were driven by residual difficulty differences, we would expect the two control groups to exhibit distinct patterns of behavior — for example, one group aligning with and the other contradicting the trend of differences, or at least differing significantly in the magnitude of the observed effects. Such outcomes would manifest as significant interactions between the between-subject factor control condition and the within-subject factor task. However, our results showed no such interactions across the three types of identified clusters. Participants from both control groups exhibited similar patterns, with both showing strong and consistent differences between the two task conditions, in either power of coupling comparison (Fig. S2).

Oscillatory coupling has been proposed to mediate communication between remote areas by providing dynamic windows of phase relations optimally geared to permit signal transmission(Engel et al., 2001; Engel et al., 2013; Fries, 2005, 2015; Womelsdorf et al., 2007). This notion implies that behavioral performance should also be associated with changes in the phase relations in the network and more efficient at particular phases of coupling, but impaired at other phase relations between the recorded sites (Fries, 2015; Rohenkohl et al., 2018). We examined how the behavioral performance was tuned by the theta phase in the prediction-related network identified in our cluster analysis. In the prediction condition, both accuracy and response times exhibited a sinusoidal tuning by the theta phase in the prediction phase of the trials. This effect was much weaker in the memory trials. In the prediction task, the participants needed to acquire the information on the sequence and, thus phase relations may have been more strongly employed to coordinate signals across the involved brain areas. In contrast, the memory task possibly did not demand the same amount of coordination across regions of the prediction network. Similar effects have been demonstrated in cellular recordings in animal studies (Ni et al., 2016; Womelsdorf et al., 2007). A recent study showed that when phase relation between V1 and V4 neurons was optimal, monkeys tended to perform faster, during a visual attention task (Rohenkohl et al., 2018). Our study is, to our best knowledge, the first demonstration of this sinusoidal tuning of behavior by the phase of oscillatory coupling at the level of whole-brain recordings. Overall, our results strongly support the notion that the phase of neural coupling is of key relevance for coordination of the information flow between separate brain regions (Engel et al., 2013; Fries, 2005).

In a serial prediction task, preceding stimuli should act as priors for processing of the current item in the sequence which, in turn, should influence the processing of subsequent stimuli. We therefore investigated whether power changes occurring during the prediction task also reflected the stimulus context, i.e., whether the preceding stimulus would have a contextual effect on the current one, and whether processing of the current item already predicted processing of a subsequent item. Both were observed to be the case and to be reflected in changes of beta power, which was larger if the same item appeared before or after the current item, suggesting that the power difference related to differences in the expectation and storage of information in the network. The storage effect was more pronounced during the prediction than during the sampling phase, possibly reflecting that participants needed to actively access the remembered information of previous items to update the prediction of the forthcoming events. The expectation effect could only be observed in the prediction phase – only when they knew the sequence. Interestingly, this effect mainly involved the motor, supplementary motor, and premotor regions. This strongly supports the idea that prediction is an active process, which relies on areas involved in action control even in the absence of an overt motor response (Bubic et al., 2010; Engel et al., 2001). The results of our phase-amplitude coupling analysis for the theta and beta signals in the prediction network suggest that beta power, through coupling to the local theta phase, might dynamically modulate the information transfer through the long-range theta phase-phase coupling. Similar phase-amplitude coupling between theta and beta signals has been observed in previous studies, e.g., in visual working memory paradigms (Daume et al., 2017).

Taken together, the power and coupling changes observed for the prediction-related network suggest the following possible scenario underlying the processing of sequence information in our task setting: In the sampling phase, the encoding of the presented items may be differentially modulated by changes in beta power already in sensory regions, depending on the task (prediction vs. memory) and on the contextual information. In the prediction phase of the trials, incoming new information is compared with the stored information on the sequence, which induces further modulation of local beta power. Local phase-amplitude coupling between beta and theta signals may then shape the transfer of the task-relevant information by long-range theta band coupling in the prediction network. After the transfer of prediction-related information, through theta phase coupling, to target motor-related areas, the information might be recoded again through theta phase to beta power coupling, which then might have an impact on response selection. By demonstrating the involvement of several frequency bands in the multisensory prediction task studied here, our results support recent concepts on multi-timescale information processing which emphasize the need to deploy different time windows for segregation and integration of information (Senkowski & Engel, 2024; Wolff et al., 2022).

In conclusion, we have investigated the neural mechanisms of multisensory sequence prediction. Our data provide direct evidence that oscillations and phase coupling across remote regions play an important role in prediction, and might serve as a general mechanism for cross-regional communication and collaboration. In particular, our results support the notion that the phase of oscillatory signals can sinusoidally modulate behavioral performance. Furthermore, our data underscore the relevance of cross-frequency coupling for establishing multi-timescale processing dynamics. These findings substantially corroborate and extend previous studies on the functional role of coupled oscillations in large-scale communication in the brain.

## 5. Ending sections

### 5.1 Data and Code Availability

The code used for the analysis in this study is publicly available in the following GitHub repository: https://github.com/neuro-science/neuro_data_analysis.

The datasets supporting all figures in this paper can be accessed and downloaded via the following link: https://box.uke.de/s/zHxJRiANYTWnOx2.

### 5.2 Author Contributions

P. W. : Conceptualization, Data Curation, Formal Analysis, Investigation, Methodology, Project Administration, Resources, Software, Validation, Visualization, Writing – Original Draft Preparation, Writing – Review & Editing; A. M. : Conceptualization, Writing – Review & Editing; J. D. : Conceptualization, Writing – Review & Editing; G. X. : Writing – Review & Editing; A. K. E. : Conceptualization, Funding Acquisition, Methodology, Project Administration, Resources, Supervision, Writing – Review & Editing

### 5.3 Funding

A.K.E. acknowledges support for this work from the DFG (SFB936-178316478-A3; TRR169-261402652-B1/B4/Z2) and from the European Union (project cICMs, ERC-2022-AdG-101097402). Views and opinions expressed in this article are those of the authors only and do not necessarily reflect those of the European Union or the European Research Council. Neither the European Union nor the granting authority can be held responsible for them.

## Acknowledgements

We thank Guido Nolte for methodological discussions and Till Schneider for valuable comments on the experimental design. We thank Roger Zimmermann, Christiane Reissmann, and Karin Reimann for help with MEG data collection and participant recruitment.

---

**Fig. S1.**
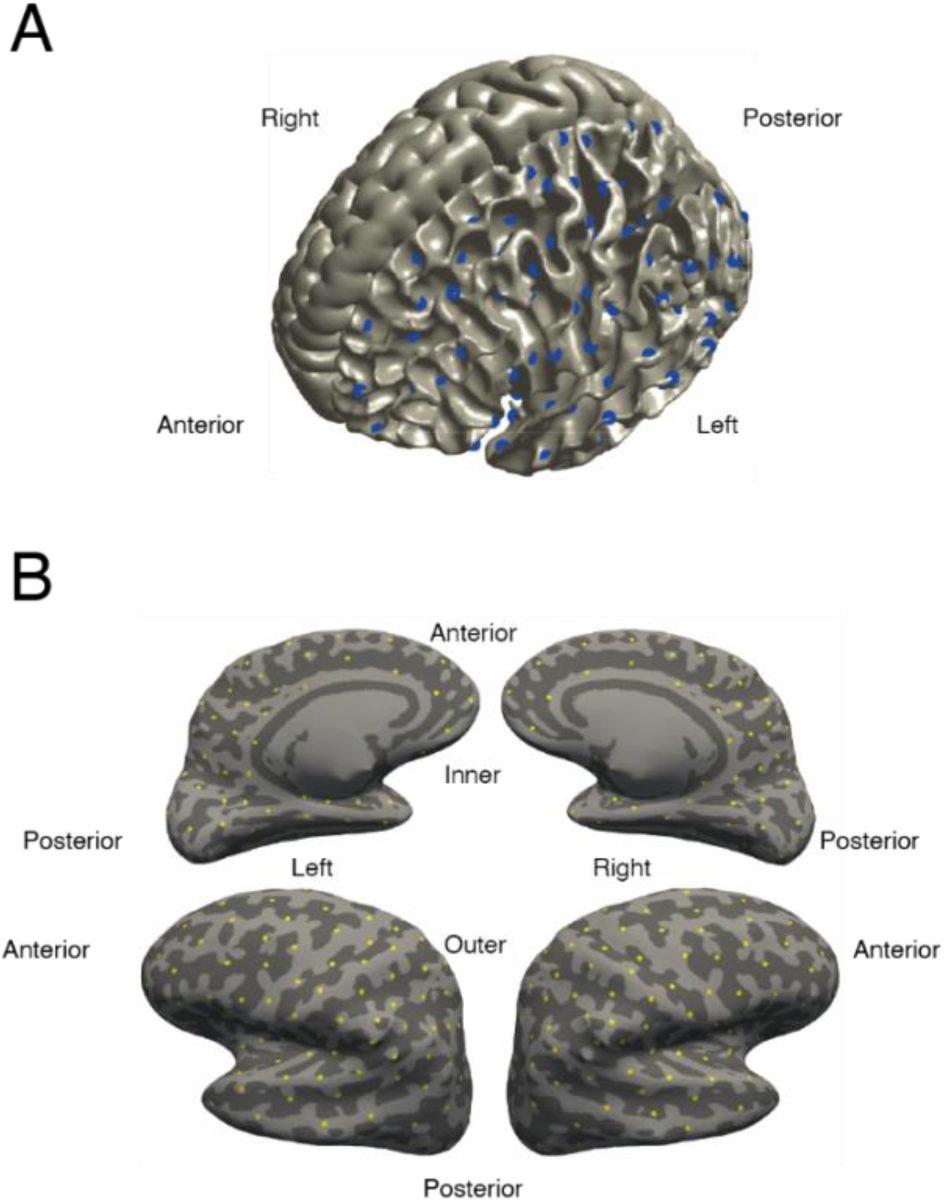
Definition of the source space. (A) White matter (left hemisphere) and pial (right hemisphere) surfaces of the template brain. Source locations were evenly distributed on the surface of the white matter, as indicated by the blue dots in the left hemisphere. (B) The inflated view of the template brain. Source locations are indicated by the yellow dots on the inflated surface.

**Fig. S2.**
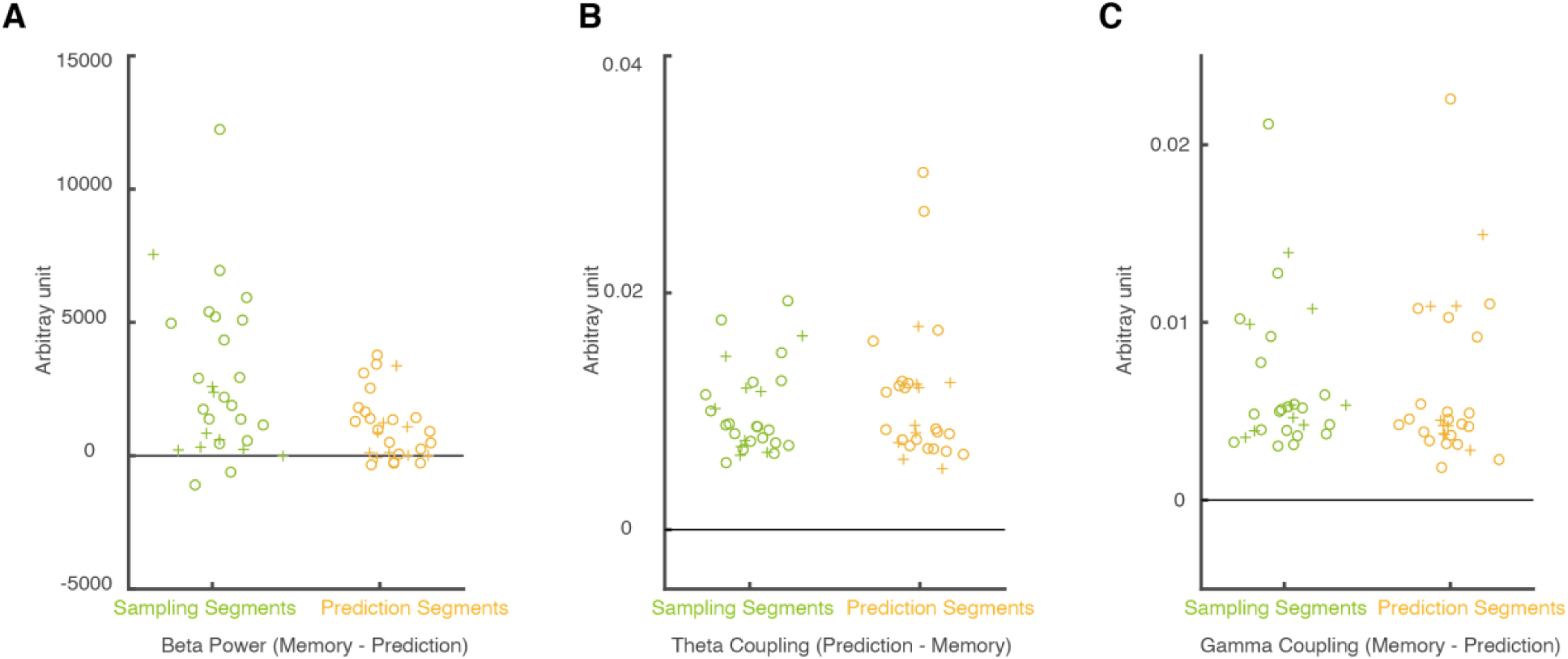
Between-task differences in the identified clusters across control conditions. The panels show differences between the prediction and memory tasks for (A) beta power (memory – prediction), (B) theta coupling (prediction – memory), and (C) gamma coupling (memory – prediction). The subtraction order was chosen to ensure positive values, facilitating easier comparison across different measures. Results are shown separately for sampling segments and prediction segments. Each symbol represents a participant, with two different control conditions. Horizontal black lines indicate zero differences.

**Fig. S3.**
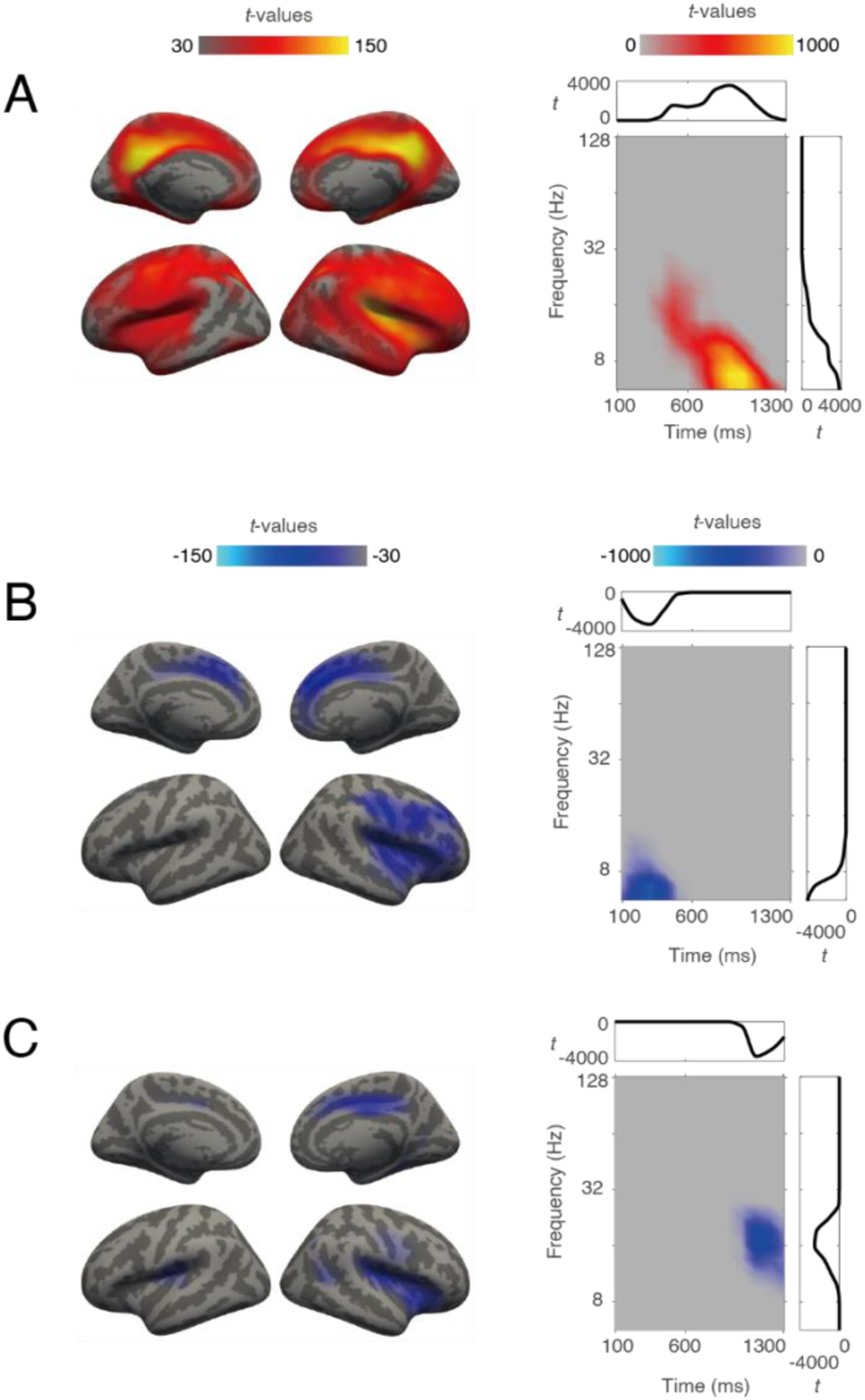
Cluster analysis of power during response segments. (A - C). In each panel, both spatial (left) and time-frequency (right) distribution are illustrated. Color indicates accumulated *t* values. Side plots in the box are integrated *t* values against time (top, x-axis: time in ms, y-axis: *t* value) or frequency (right, x-axis: *t* value, y-axis: frequency in Hz).

**Fig. S4.**
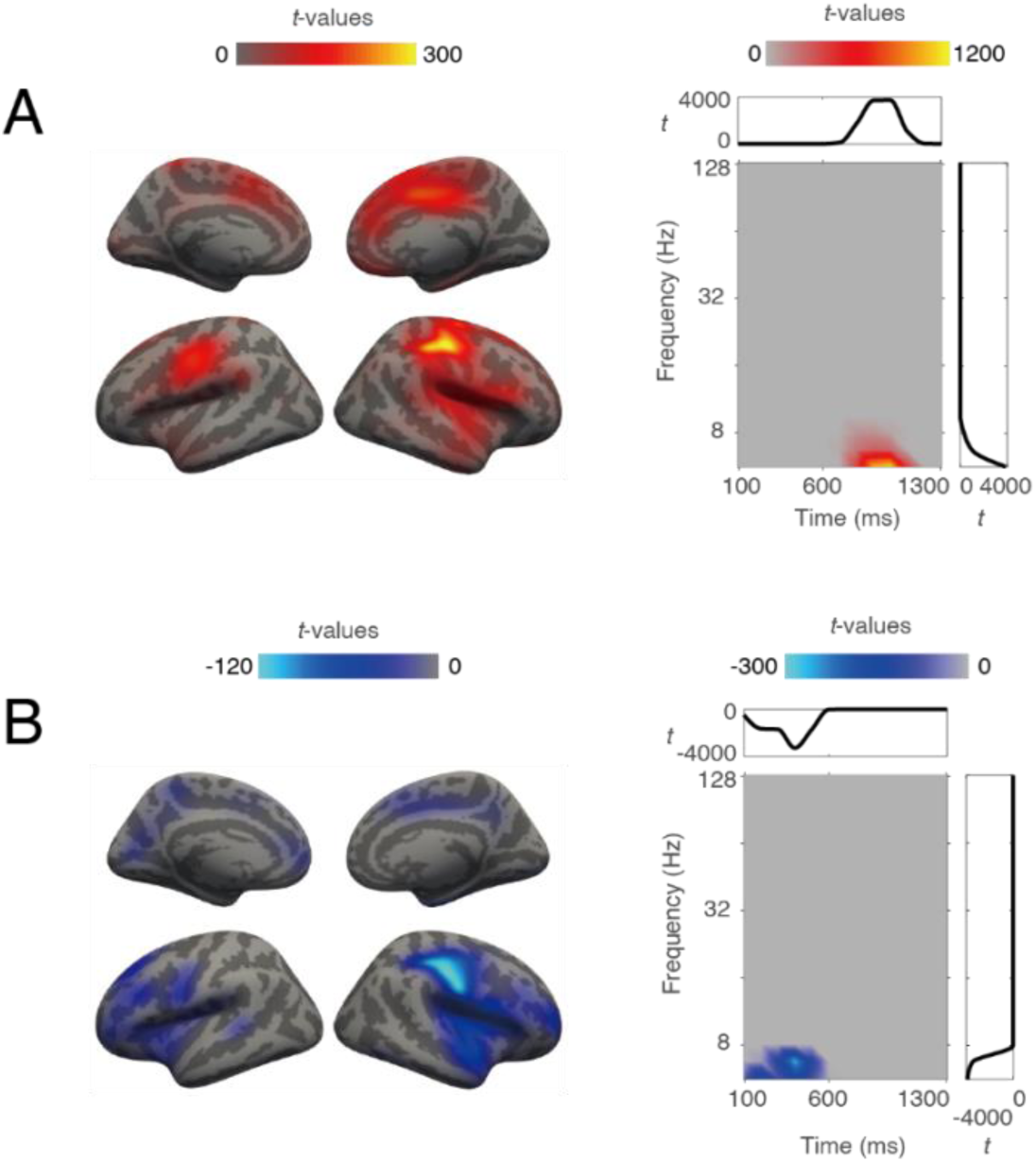
Cluster analysis of coupling during response segments. In each panel, both spatial (left) and time-frequency (right) distribution are illustrated. Color indicates accumulated *t* values. Side plots in the box are integrated *t* values against time (top, x-axis: time in ms, y-axis: *t* value) or frequency (right, x-axis: *t* value, y-axis: frequency in Hz).

